# Age-associated elevated inflammation and immune pathways in the mice heart are linked to Adiponectin-Adipor1-RelA signaling

**DOI:** 10.1101/2023.04.28.538681

**Authors:** Tinku Gupta, Akash Gujaral, Shivanshu Chandan

## Abstract

Inflammatory gene profiles using RNA seq analysis were studied by measuring pro-inflammatory cytokines and chemokines levels. qRT-PCR and Western blot analysis were used to validate the expression profile of these inflammatory mediators. Using flow cytometry analysis, CD11b+ monocytes and CD64+ Ly6C were quantified in the young and old hearts. The inflammatory response, Adipor1 and Adipor2 gene expression, RelA nuclear translocation and the effects of adiponectin in LPS-stimulated or Adipor1 silenced H9C2 cells were studied. Gene ontology analysis using differentially expressed genes revealed an enrichment of immune response pathways in the old mice hearts when compared to young mice hearts. Western blot analysis confirmed the down regulation of several anti-inflammatory proteins and the upregulation of pro-inflammatory proteins including CD68, NF-kB1 and Rel-A, in the old mice hearts. Flow cytometry suggested an infiltration of CD11b+monocytes and CD64+ Ly6C-high macrophages in the old mice hearts compared to younger hearts confirming an increased inflammation in the older hearts. Mechanistically, to understand if the Adiponectin-Adipor1-NFkB axis regulates inflammation in the aging heart, Adipor1 and Adipor2 genes were silenced in H9c2 cardiomyocytes. Immune response genes were elevated in the Adipor1 silenced H9c2 cells but not in Adipor2 silenced cells. Pretreatment with Adiponectin (APN) attenuated the Adipor1 silenced or lipopolysaccharides (LPS)-stimulated expression of inflammatory genes in H9c2 cardiomyocytes. APN also attenuated the nuclear translocation of RelA and induction of immune response genes in Adipor1 silenced or LPS-challenged H9c2 cardiomyocytes. APN-AdipoR1-RelA signaling might be a novel therapeutic target for the treatment of inflamed elderly hearts.

## 1. Introduction

Adiponectin is a hormone produced by adipocytes cells that store fat [1, 2]. Adiponectin has various physiological effects, including regulation of glucose metabolism and insulin sensitivity and protection against inflammation. Adiponectin binds to two receptors on the cell surface, AdipoR1, and AdipoR2, to activate signaling pathways that promote its effects [3, 4].

One of the critical signaling pathways activated by adiponectin is the NFkB pathway^3, 4, 5, 6^. NF-kB is a transcription factor that regulates gene expression and promotes inflammation. In the aging heart, adiponectin-AdipoR1-NF-kB signaling plays an important role in maintaining heart function. Studies have shown that reduced adiponectin levels are associated with an increased risk of cardiovascular disease, including heart failure, in the elderly [5, 6, 7].

Adiponectin has been shown to improve heart function by reducing oxidative stress and inflammation, promoting angiogenesis (formation of new blood vessels), and regulating glucose and lipid metabolism [8, 9]. Adiponectin also stimulates the production of nitric oxide, a vasodilator that helps maintain blood flow to the heart [10].

In addition, adiponectin has been shown to inhibit the activation of NFkB, which is thought to contribute to the decline in heart function during aging. By inhibiting NFkB activation, adiponectin helps to reduce oxidative stress, inflammation, and apoptosis (programmed cell death) in the aging heart [5–10].

The adiponectin-AdipoR1-NFkB signaling is thus crucial in maintaining heart function in the elderly population. Further research is needed to understand the underlying mechanisms and develop therapeutic strategies to enhance adiponectin signaling in the aging heart.

Given that the regulators of cardiac inflammation have not been described extensively and rarely have been correlated with cardiac adiponectin-AdipoR1-NFkB signaling directly, we sought to identify critical regulators of cardiac inflammation and their association with this signaling using young and old mice hearts.

Using RNA-seq profiles of the left ventricular (LV) tissue of young and old mice hearts, we performed a screen of genes regulating cardiac inflammation. We identified several pro-inflammatory and anti-inflammatory genes, including Adipor1, Adipor2, and RelA, that belong to diverse immune response pathways in the aging heart. Using Adipor1 silenced cardiomyocytes, we provided evidence that suggests links between Adiponectin-Adipr1-RelA signaling and immune response in the older heart. We thus conclude that the upregulation of specific inflammatory and immune response genes in older mice hearts might be linked to the imbalanced Adiponectin-AdipoR1-RelA axis. APN-AdipoR1-RelA signaling might be a novel therapeutic target for the treatment of inflamed elderly hearts.

## 2. Results

### 2.1) Global reprogramming of immunological changes, especially inflammatory cytokines and chemokines signaling in the left ventricular tissue of the old mice

Inflammation is widely recognized to have a role in the pathogenesis of various diseases, including cardiovascular disease associated with aging [2, 11, 12]. To define the immunological changes in the aging heart, we performed RNA sequencing (RNA-seq) using left ventricular (LV) tissue of the heart of 3 months (3M) and 18 months (18M) old mice. Using a 1.3-fold cutoff and false discovery rate (FDR) <0.05 threshold for inclusion, we identified approximately 2329 and 1718 differentially expressed genes in LV tissue of mice at 3M and 18M of age, respectively. Venn diagram revealed ∼1588 uniquely expressed genes and ∼977 uniquely expressed genes in LV tissue of mice at 3M and 18M, respectively (Fig. 1A). The top ten enriched Reactome pathways observed in the heart of 3M old mice were related to the citric acid cycle, pyruvate metabolism, mitochondrial biogenesis, and inactivation of G-CSF signaling (Fig. 1B, Supplementary Table 1). Gene ontology analysis (top 10 Reactome pathways) using differentially expressed genes indicated an enrichment of immunological pathways especially inflammatory response, Complement activation, and immune response pathway in older mice hearts (Fig. 1C, Supplementary Table 1).

**Figure 1.**
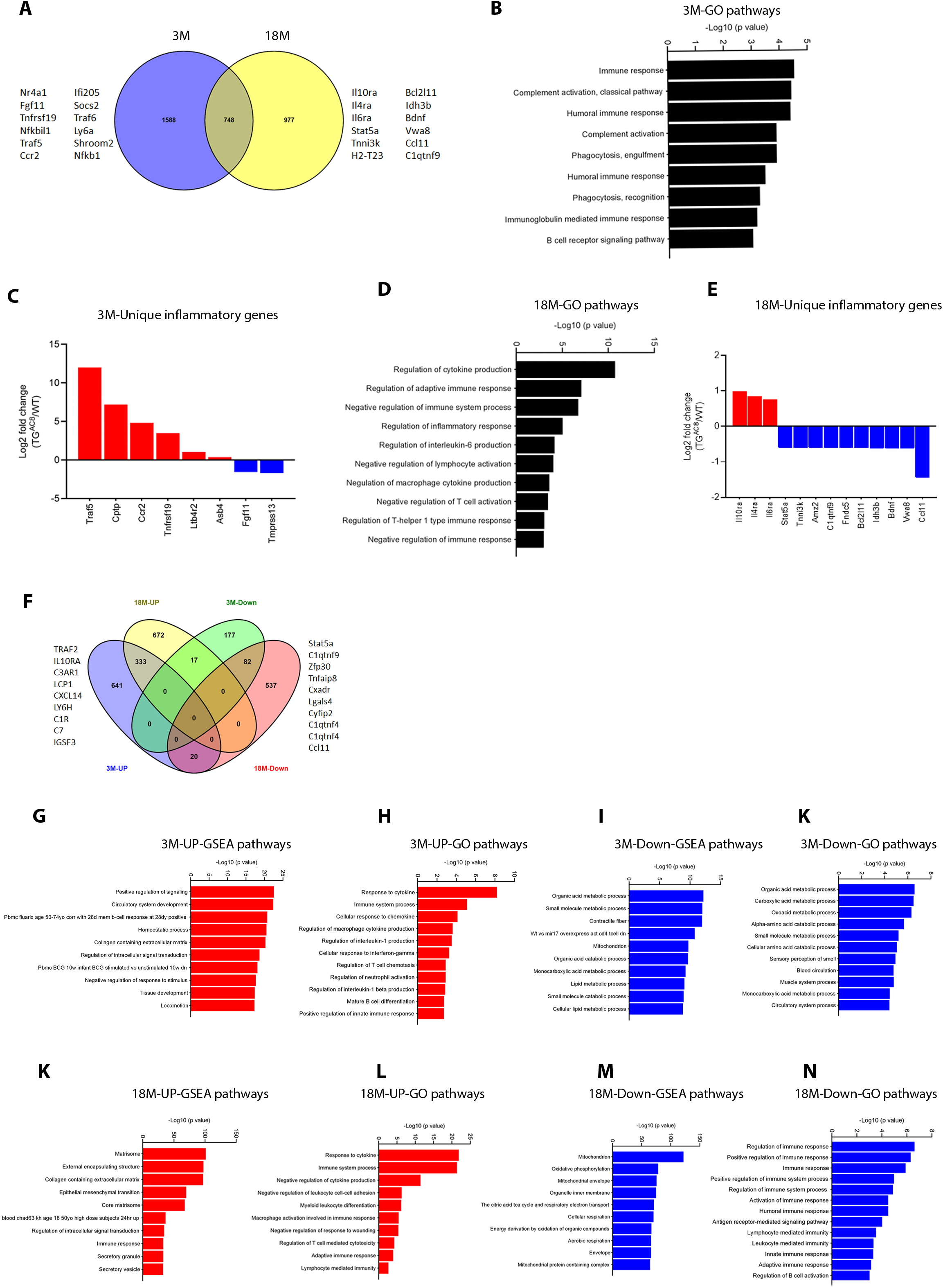
Global reprogramming of immunological changes, especially inflammatory cytokines, and chemokines in the left ventricular tissue of the heart of old (18M) mice. **(A)** Venn diagram was performed using differentially expressed genes identified in RNA-seq (the cutoff for fold change >1.35, and *P*<0.05). The Venn diagram represents the common and uniquely identified genes between young and old mice hearts. **(B), (C)** Gene ontology analysis for top enriched Reactome pathways was performed using differentially expressed genes (the cutoff for fold change >1.35, and *P*<0.05) identified between young and old mice hearts. **(D)** Heatmap was performed via online software Heatmapper using differentially expressed genes (the cutoff for fold change >1.35, and *P*<0.05) between young and old mice hearts. The heatmap represents the specific inflammatory genes, including cytokines and chemokines, differentially expressed between young and old mice hearts.

We then analyzed the specific inflammatory genes differentially expressed in 3M and 18M mice hearts. Of the top differentially expressed immune response and inflammatory genes, Adipor1, Adipor2, Socs2, Socs3, Il10ra, Il13ra1, Fabp4, Il12rb2, and Arg-1 were downregulated, and CD68, Cd163, RelA, NF-kB1, Leprotl1, Cx3cr1, Cx3cl1, Ccl11 and IL6ra were upregulated in the old mice hearts when compared to the young mice hearts (Fig.1D). To understand the cardiac function, we observed via echocardiography that the old mice heart had significantly (*P*<0.001) lower ejection fraction than young mice (Supplementary figure A, B). These results suggest a possible activation of immune response signaling, decreased Adiponectin-Adipor1 or Adipor2 signaling, and an imbalanced pro-inflammatory and anti-inflammatory signaling in the old (18M) mice hearts compared to younger (3M) mice hearts.

### 2.2) Elevated myocardial inflammation and infiltration of immune cells in the aging heart

Immune cell infiltration and aggressive cytokine response contribute to the pathogenesis of an aging heart. In addition, immune cells, especially cardiac resident macrophages, facilitate the heart’s electrical conduction by coupling to cardiomyocytes in the atrioventricular node [7, 18, 19]. We identified several pro-inflammatory and anti-inflammatory molecules in our RNA-seq data (Supplementary Table 1). We validated these changes using the total LV tissue lysates via Western blot analysis. Notably, the pro-inflammatory proteins such as CD68 (macrophage marker), NF-kB1, and RelA were upregulated and the anti-inflammatory proteins including Adipor1, Adipor2, Socs2 (suppressor of cytokine signaling type2), and Socs3 (suppressor of cytokine signaling type 3) were downregulated in the LV tissues of 18M old mice compared to 3M old mice (Fig. 2A). We then confirmed the nuclear translocation of RelA, a subunit of NF-kB and master regulator of inflammation was using a nuclear lysate of LV tissue of 3M and 18M old mice (Fig. 2B). To assess if inflammatory response accompanies with the elevated infiltration of immune cells in the aging heart, using flow cytometry analysis, we quantified the CD45+ CD11b+ monocytes, CD64-Ly6c+ monocytes and CD64+ Ly6c-high macrophages in the heart. Flow cytometry analysis confirmed the elevated infiltration of these cells in the old heart when compared to the young heart (Fig. 2C, 2D). These results confirmed the elevated myocardial inflammation in the aging heart.

**Figure 2.**
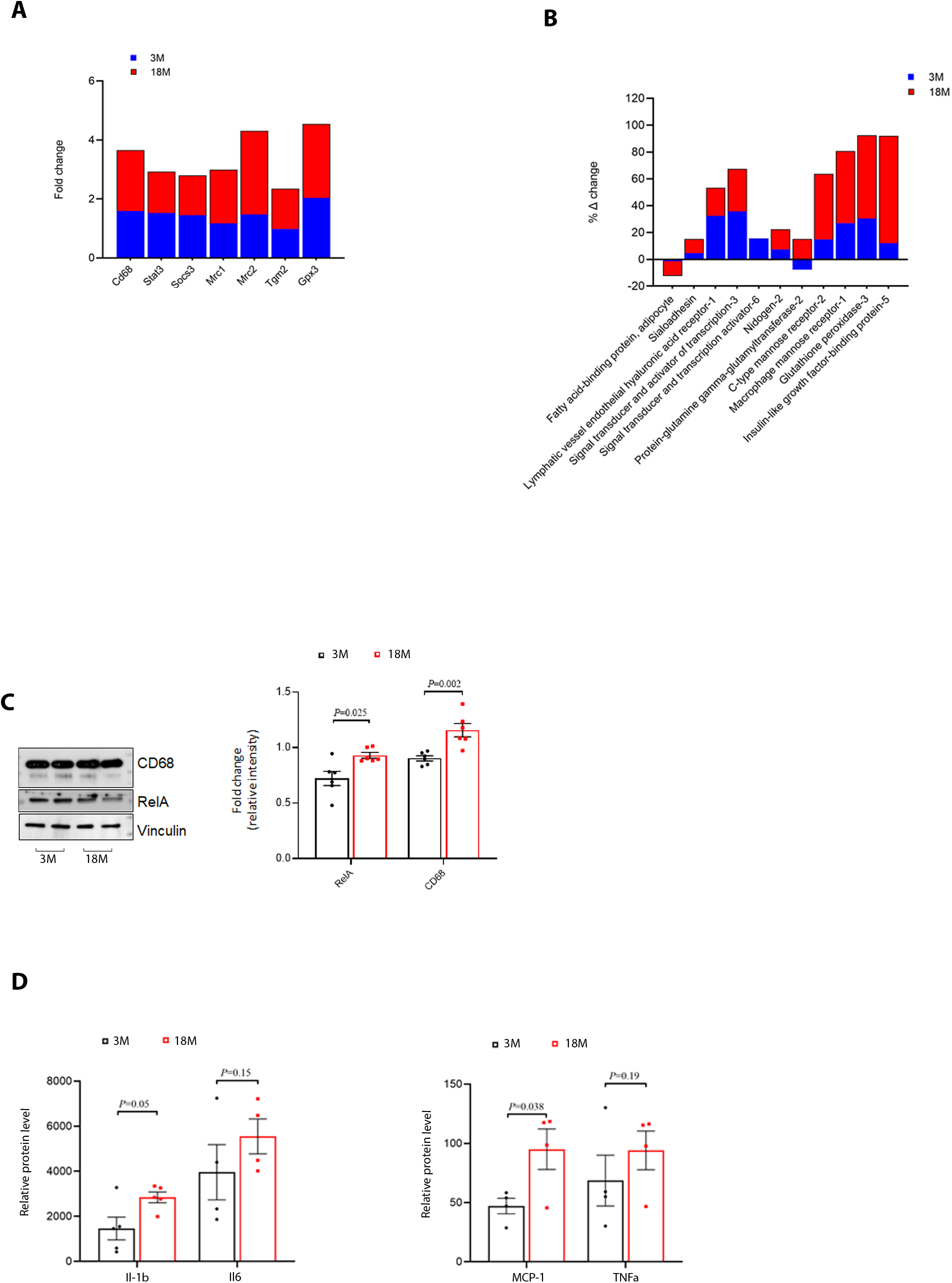
Elevated myocardial inflammation and infiltration of immune cells in the aging mice heart. **(A)** Representative Western blot analysis using total LV tissue lysates of young (3M) and old mice (18M) (n=4 per group). Validation of expression of the pro-inflammatory proteins and the anti-inflammatory proteins between LV tissue of 3M and 18M old mice **(B)** Representative Western blot analysis using a nuclear lysate of LV tissue of 3M and 18M old mice (n=4 per group). **(C), (D)** Representative FACS plot of CD45+ CD11b+ monocytes, CD64-Ly6c+ monocytes, and CD64+ Ly6c-high macrophages between LV tissue of 3M and 18M old mice. Flow cytometry analysis revealed elevated myocardial inflammation in the aging mice heart compared to young mice hearts.

### 2.3) Adipor1 but not Adipor2 silencing induces the nuclear translocation of RelA and the expression of immune response genes in H9c2 cardiomyocytes

Adipor1 is an antiinflammatory chemokine that regulates immune response and inflammation. To understand if the immunological changes observed in aging hearts are linked to the Adipor1/Adipor2-RelA signaling, we silenced Adipor1 or Adipor2 gene in H9c2 cardiomyocytes. Immune response genes, including IL-1□, IL6, and TNF-α were upregulated, and the anti-inflammatory factor Arg-1 gene was downregulated in Adipor1-silenced H9c2 cardiomyocytes. AdipoR1 silencing also stimulated the nuclear translocation of the RelA protein, suggesting that RelA might have increased the transcription of immune response genes in these cells (Fig. 3A-3F). Adipor2 silencing did not significantly change the expression of IL-1□, IL6, and TNF-α genes and down regulated the Arg-1 (anti-inflammatory gene) gene expression (Fig. 3G-3K). These results suggest that Adipor1 but not Adipor2 might regulate the expression of inflammatory response genes in H9c2 cardiomyocytes.

**Figure 3.**
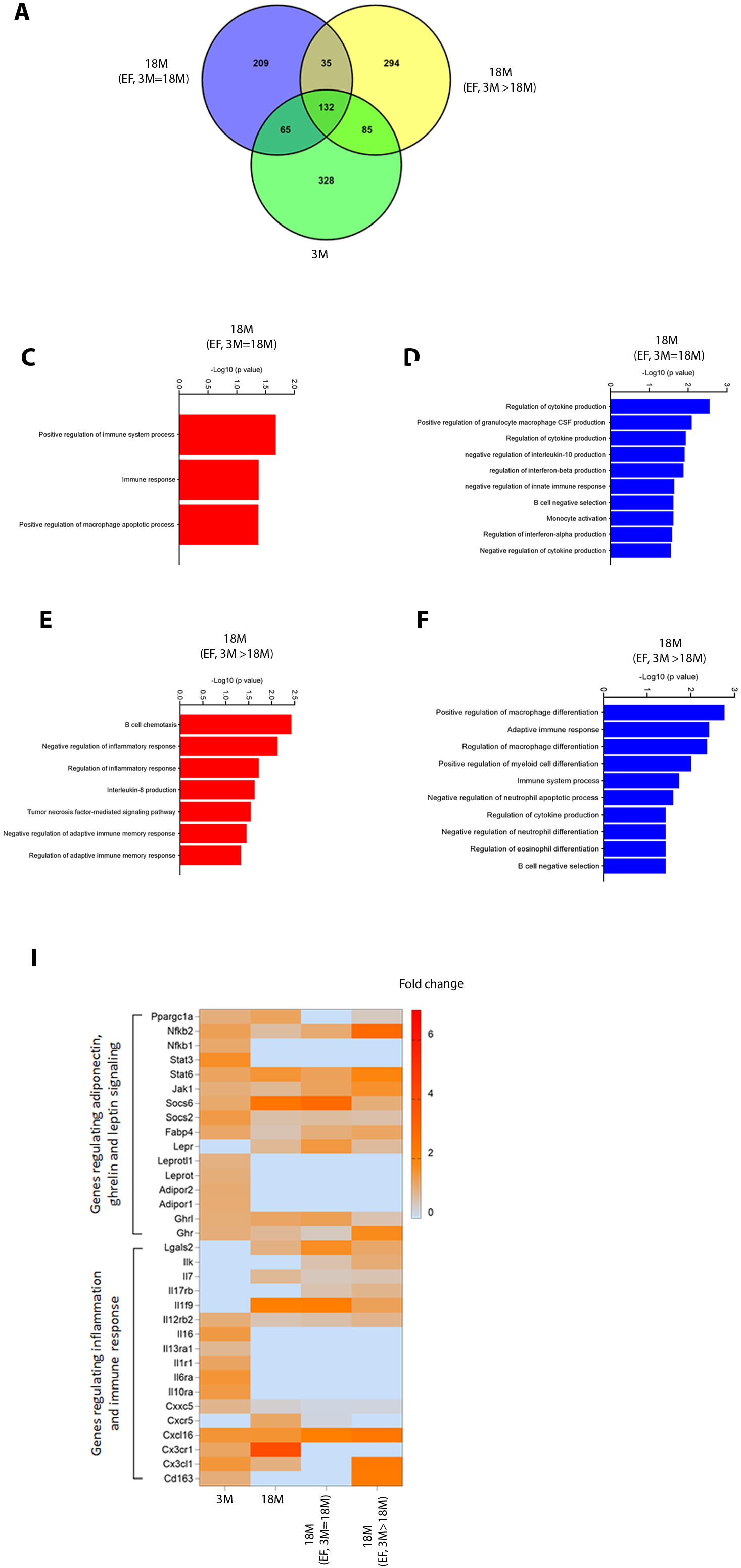
Adipor1 but not Adipor2 silencing induces the nuclear translocation of RelA and the expression of immune response genes in H9c2 cardiomyocytes. **(A)** Representative Western blot analysis of Adipor1 protein using whole cell lysate of H9c2 cardiomyocytes transfected with either control vector or si-Adipor1 (n=3 per group). H9c2 cells were silenced using shAdipor1 plasmid for 24 hours and harvested for gene and protein expression analysis. **(B)-(E)** qRT-PCR analysis of inflammatory genes (IL-1□, IL6, TNFα and Arg-1) using whole cell lysate of H9c2 cardiomyocytes transfected with either control vector or si-Adipor1 (n=6 per group). **(F)** Representative Western blot analysis of RelA protein using a nuclear lysate of H9c2 cardiomyocytes transfected with either control vector or si-Adipor1 (n=3 per group). **(G)** Representative Western blot analysis of Adipor2 protein using whole cell lysate of H9c2 cardiomyocytes transfected with either control vector or si-Adipor2 (n=3 per group). H9c2 cells were silenced using siAdipor2 plasmid for 24 hours and harvested for gene and protein expression analysis. **(H)-(K)** qRT-PCR analysis of inflammatory genes (IL-1□, IL6, TNFα and Arg-1) using whole cell lysate of H9c2 cardiomyocytes transfected with either control vector or si-Adipor1 (n=6 per group).

### 2.4) Adiponectin (APN) attenuates the induced expression of immune response genes and nuclear translocation of RelA in LPS-induced or Adipor1 silenced H9c2 cardiomyocytes

Adipokines decrease the levels of chronic inflammatory state cytokines and thereby contribute to the pathogenesis of cardiovascular diseases [2–10]. We observed decreased level of adiponectin in the LV tissue of aging mice when compared to young mice (Supplementary figure C, D). Interestingly, leptin, an inflammatory chemokine level, was higher in the LV tissue of old mice. We asked if APN supplementation attenuates the expression of inflammatory cytokines and nuclear translocation of RelA in Adipor1 silenced or LPS-induced H9c2 cardiomyocytes. LPS stimulation upregulated the expression of immune response genes and nuclear translocation of RelA in H9c2 cardiomyocytes. APN supplementation (10μg/ml) attenuated the induction in the expression of immune response genes (IL-1□, IL6, and TNF-α) and nuclear translocation of RelA protein in LPS-stimulated H9c2 cardiomyocytes (Fig. 4A-4E). To investigate if APN via Adipor1-RelA signaling increases the expression of immune response genes in the heart, we silenced the Adipor1 gene via siRNA in H9c2 cardiomyocytes and supplemented the cells with APN. APN supplementation attenuated the induction in the expression of immune response genes, and nuclear translocation of RelA protein in Adipor1 silenced H9c2 cardiomyocytes (Fig. 4F-4I). These results suggest that APN via Adipor1-RelA might regulate the expression of inflammatory response genes in H9c2 cardiomyocytes.

**Figure 4.**
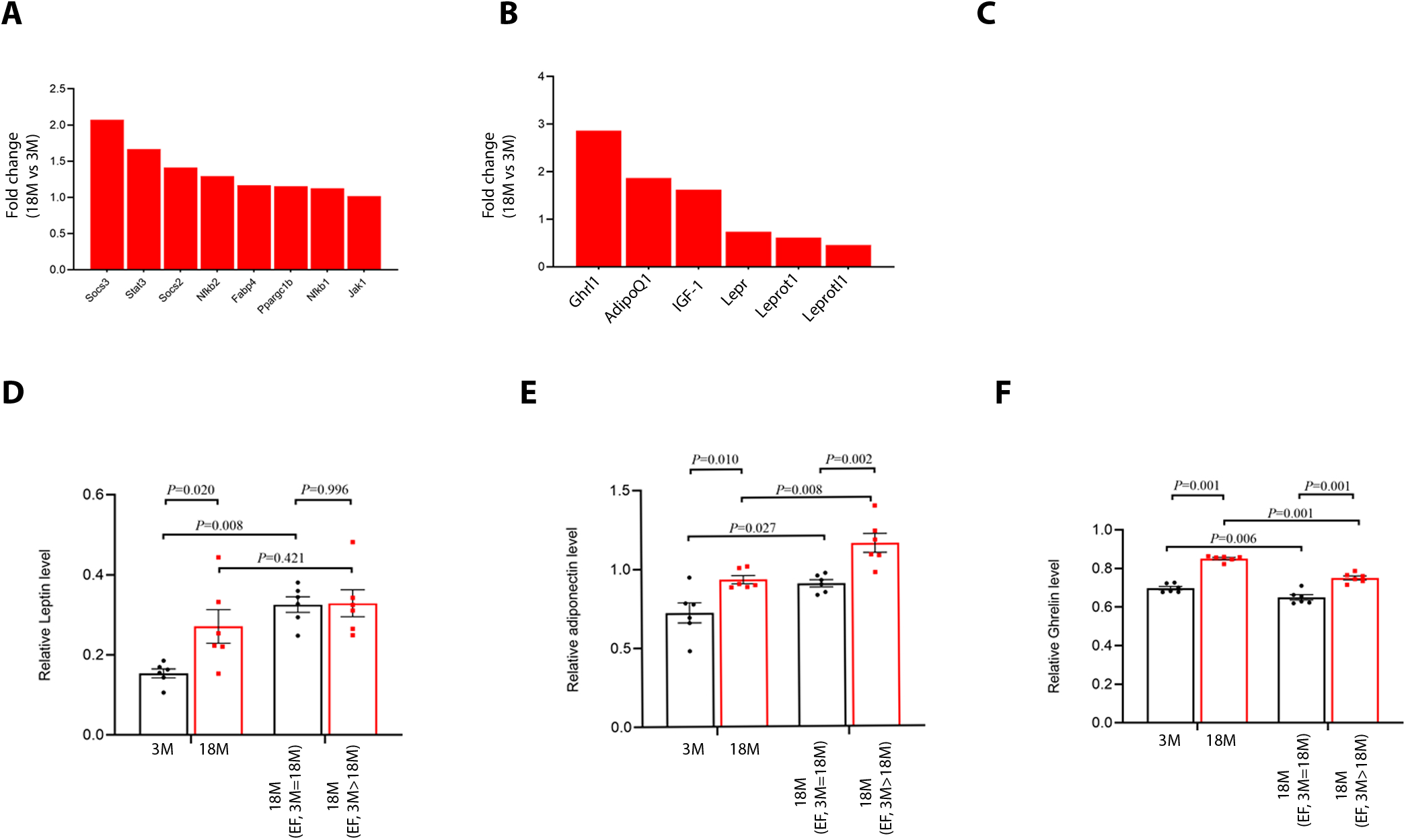
Adiponectin (APN) attenuates the induced expression of immune response genes, and nuclear translocation of RelA in LPS-induced or Adipor1 silenced H9c2 cardiomyocytes. **(A)-(D)** qRT-PCR analysis of inflammatory genes (IL-1□, IL6, TNFα and Arg-1) using whole cell lysate of H9c2 cardiomyocytes pretreated with APN and then challenged with LPS (n=6 per group). **(E)** Representative Western blot analysis of RelA protein using a nuclear lysate of H9c2 cardiomyocytes (pretreated with APN and then stimulated with LPS) (n=4 per group). **(F)** Representative Western blot analysis of Rela protein using a nuclear lysate of H9c2 cardiomyocytes pretreated with APN and then transfected with either control vector or si-Adipor1 (n=3 per group). H9c2 cells were pretreated with APN and then silenced using siAdipor1 plasmid for 24 hours and harvested for gene and protein expression analysis. **(G) - (I)** qRT-PCR analysis of inflammatory genes (IL-1□, IL6, and Arg-1) using whole cell lysate of H9c2 cardiomyocytes pretreated with APN and then transfected with either control vector or si-Adipor1 (n=6 per group).

## 3. Discussion

Activation of the immune system and inflammatory gene program contributes to pathological changes in the aging heart [13–15]. Immune response and inflammatory changes are among the several adaptive changes in the heart in response to diverse physiological stresses and pathological injury. The immune system’s homeostatic mechanisms decline with advancing age and thus can increase susceptibility to transition from adaptive remodeling to a failing heart^15^. Dissecting the inflammatory changes associated with aging could aid in identifying novel therapies for aging-associated cardiac failure.

We have found evidence for the potential involvement of Adiponectin (APN)-Adipor1-RelA signaling in the inflammatory gene program of the aging heart. The role of this signaling axis on cardiac inflammatory response has not been completely understood [2, 15]. Our study has identified several pro-inflammatory and anti-inflammatory genes associated with several immune response pathways in the aging heart. We then observed the association between activation of the inflammatory gene program and reduced Adipor1 expression and increased nuclear translocation of RelA, a master regulator of inflammation in old mice hearts. Additionally, we provided evidence that APN supplementation attenuated the cardiac inflammatory response in Adipor1 silenced and LPS-stimulated H9c2 cardiomyocytes. We conclude that the Adiponectin via Adipor1-Rela signaling regulates the inflammatory response in the H9c2 cardiomyocytes.

Inflammation, especially immune cells, infiltrate the heart, modulates its contractile function, produces soluble factors, including cytokines and chemokines during acute inflammation, phagocytes apoptotic cells, and repairs the scar left after myocardial infarction ischemia [16, 17, 18]. A coordinated immune system is required for the steady-state function of the heart [15]. A constant decline in immune system function has been observed in the aging heart, which hinders the ability of the heart to adapt to various physiological and pathological stress conditions, including heart failure. Aging is associated with the infiltration of myeloid cells, especially macrophages in the heart, which suppress inflammation-mediated immune-senescence [14, 15, 19]. We provide evidence of reprogramming in the inflammatory process via RNA-seq analysis in the old mice’s heart. A unique set of pro-inflammatory genes, especially RelA and CD68, and immune response pathways are upregulated in the old mice heart, and Adipopr1, Adipor2, and Socs2 genes, the anti-inflammatory factors, are down regulated in these older mice hearts. We also observed an infiltration of immune cells, especially CD11b+ monocytes, and Ly6c-high macrophages in the heart of old mice, as observed in earlier studies.

Adipor1 silencing increases the pro-inflammatory cytokine expression in BV2 and microglial cells [3]. The inflammatory response was specific to the Adipor1 receptor only but not Adipor2 receptor. Decreased expression of Adipor1 expression is also observed in the heart of older mice. In concordance with this study, we observed a decreased expression of Adipor1 receptors in the aging mice heart. Adipor1 silenced H9c2 cardiomyocytes had a lower inflammatory response confirming the link between Adipor1 and inflammatory response. A similar relationship was not seen between Adipor2 and inflammatory response. Adipokine, especially adiponectin, is an anti-inflammatory mediator emerging as a therapeutic target for regulating inflammation and immune responses in several chronic disease conditions such as obesity, diabetes mellitus, and cardiovascular diseases [2, 3]. Adiponectin, an adipose tissue-derived plasma protein, regulates glucose levels, lipid metabolism, and insulin sensitivity [20]. Elevated plasma adiponectin is associated with obesity-induced endothelial dysfunction and hypertension and protects against atherosclerosis, myocardial infarction, and diabetic cardiomyopathy [9, 10]. Lack of adiponectin is associated with the increased endoplasmic reticulum (ER) stress, increased oxidative stress, increased pro-inflammatory responses, and decreased protective cytokines in the heart [11,13]. We sought to quantify the adiponectin level in the aging heart and found a lower adiponectin level in older mice. We next asked if adiponectin supplementation reversed or attenuated the inflammatory response in the LPS-challenged or Adipor1 silenced cardiomyocytes.

Interestingly, adiponectin supplementation attenuated the inflammatory response, increased the anti-inflammatory Arg-1 gene expression, and inhibited the nuclear translocation of RelA in Adipor1 silenced and LPS-stimulated H9c2 cardiomyocytes. These findings were in line with several earlier observations, highlighting that adiponectin can regulate the pro-and anti-inflammatory signaling and energy homeostasis in disease conditions, including cardiovascular diseases [22, 23].

In summary, we have discovered a set of inflammatory genes that contribute to inflammation in the aging heart. Our study provides an in-depth analysis of the inflammatory profile of the heart of young and old mice. It reveals that decreased anti-inflammatory gene expression and increased pro-inflammatory response in the old mice heart might be associated with decreased adiponectin level and adiponectin-Adipor1-RelA signaling.

This study, however, did not focus on the transcriptomic alterations in the inflammatory gene program and their association with heart function. Another limitation of this study is that we did not use the adiponectin or Adipor1 knockout mouse models to substantiate our findings on inflammation in the aging heart.

## 4. Materials and Methods

### 4.1 Animal study

The animal studies complied with the Guide for the Care and Use of Laboratory Animals published by the National Institutes of Health (NIH). The Animal Care and Use Committee of the Rajiv Gandhi Centre for Biotechnology approved all animal study protocols. Mice were kept in the standard 12-h light-dark cycle and fed a standard chow diet ad-libitum unless stated otherwise. All the mice used in the experiments were in the C57/BL6 background. The C57BL/6J mice were used for all the studies. Young (3 months of age (3M), n=10) and old (18 months of age (18M), n=10) male mice were used for all the experiments. All the animals were regularly monitored for body weight.

### 4.2 RNA-seq

Total RNA from the left ventricular tissue of the heart was extracted from young (3M) and old (18M) mice (n=6 each). The quality of RNAseq data was assessed using a standard NGS-QC toolkit. 75 bp single-end reads generated 30 to 40 million reads per library. Raw RNA sequencing (RNASeq) reads were aligned and, after quality trimming, were mapped to the UCSC mm10 mouse reference genome and assembled using Tophat v2.0 to generate BAM files for each sample. Cufflinks v.2.1.1 were used to calculate FPKM (Fragments per Kilobase of transcript per Million mapped reads) for each sample. Differential gene expression analysis was performed with a Cuffdiff package (Cufflinks v2.1.1). Cutoff values of fold change >1.35 and FDR<0.05 were used to select differentially expressed genes between young and old mice group comparisons. The Venn diagram and gene ontology (for Reactome pathways) analysis was performed to unravel differences between young and old groups.

### 4.3 Venn diagram, Gene Ontology (GO) and Heat-map

The Venn diagram and GO analysis were performed using the differential expressed genes between young and old mice. The Venn diagram was used to compare and find uniquely expressed genes between these groups. Gene Ontology analysis for top Reactome pathways was performed to determine molecular and biological pathways enriched between young and old mice. The top enriched Reactome pathways were represented using the p-value of these pathways. A heatmap of chemokines, chemokine receptors, cytokines, cytokine receptors, and specific inflammatory mediators was generated using the online software Hatmapper.

### 4.4 Echocardiography

After an initial assessment of basal echocardiography parameters, 20 male mice (3 months old) were divided into two groups (1). Young (3M) group (n=10), and (2). Old (18 months (18M)) group (n=10). The cardiac structure and function of mice were assessed by echocardiography. Systolic intra-ventricular septal thickness (IVSs), diastolic intraventricular septal thickness (IVSd), systolic left ventricular internal dimension (LVIDs), diastolic left ventricular internal dimension (LVIDd), systolic left ventricular posterior wall thickness (LVPWs), diastolic left ventricular posterior wall thickness (LVPWd), left ventricular ejection fraction (LVEF) and left fractional ventricular shortening (LVFS) were recorded during echo measurement. The data were analyzed using the software GraphPad prism.

### 4.5 Cell culture and drug treatment

H9c2 cardiomyocytes were purchased from ATCC, USA. The cells were cultured in Dulbecco’s modified Eagle’s medium (DMEM) supplemented with fetal bovine serum (FBS, 10%) and 1% penicillin/streptomycin. The cells were maintained at 37□°C with 5% CO2 in a humidified incubator. H9c2 cardiomyocytes were quantified, pretreated with APN (10□μg/ml) for 4□h and then stimulated with LPS (100ng/ml) for 6□h in a serum-free culture medium. The cells were imaged and observed using a bright field microscope.

### 4.6 AdipoR1 and AdipoR2 silencing

AdipoR1 and AdipoR2 genes were silenced using siRNAs vectors (Santa Cruz Biotechnology). H9c2 cardiomyocytes were quantified and cultured in a six-well tissue culture plate. At approximately 80% confluency, the complete medium was replaced with serum-free medium, cells were transfected with either non-targeting siRNA or si-Adipor1 or si-Adipor2 plasmid using lipofectamine 3000 reagents (Sigma-Aldrich, USA) as per manufacturer protocol. After 6-h incubation, the siRNA-containing medium was replaced with a complete culture medium, and cells were further cultured for 18□h before harvest and analysis. The knockdown of Adipor1 or Adipor2 was confirmed using Western blot analysis; cells with knockdown between 70-80% were used for analysis.

### 4.7 Western blot analysis

The snap-frozen left ventricular tissue of the heart collected after the sacrifice of mice was homogenized in RIPA buffer with a protease inhibitor cocktail (Roche) using a prechilled Precellys bead homogenizer. The protein concentration in each sample was determined using Bicinchoninic acid (BCA) assay. 20-30ug of protein was loaded on precast Tris-Glycine SDS-PAGE gel (BioRad) and transferred to PVDF membrane by semidry transfer. Primary antibodies were incubated overnight in 3% BSA in TBST buffer. ECL signal was detected using the Amersham imager 600 imaging system. Antibodies used for immunoblot were anti-mouse Vinculin (ab91459), anti-mouse CD68 (CST-97778), anti-mouse RelA (sc-8008, 51-0500), anti-mouse TATA-binding protein/TBP (MA1-21516), anti-mouse NF-kB1 (ab7549), anti-mouse Socs2 (ab109245), anti-mouse Socs3 (ab280884), anti-mouse Adipor2 (PA5-114166), anti-mouse Adipor1 (MA5-32249), and anti-mouse PCNA (ab92552).

### 4.8 Adiponectin measurement

Total LV tissue lysates of the heart from young and old mice (n=6-8 each) were used for quantifying the adiponectin levels using a sandwich-based ELISA kit (ab108785).

### 4.9 Quantitative real-time PCR

Total RNA was isolated using Qiagen RNA mini kit from the LV tissue of the heart of young and old mice (n=6 each) and stored at −80□ until analysis. RNA was converted into cDNA using a BioRad cDNA synthesis kit; cDNA was amplified (via a TaqMan probe designed specifically for target genes) using an Applied Biosystems real-time PCR machine. The 2^-ΔΔCT^ method was used to quantify RNA, as described earlier [24, 25, 26].

### 4.10 Statistical analysis

All data are expressed as mean with SEM. Statistical analysis of data was performed using the software GraphPad Prism 9.0. P values were calculated with either unpaired/two-way t-test or one-way ANOVA followed by multiple comparisons with Tukey’s or Bonferroni test wherever applicable. P <0.05 was considered significant in all cases after corrections were made for multiple pairwise comparisons.

## Acknowledgments

We thank the Rajiv Gandhi center for Biotechnology for providing the facilities and funding the study.

## Figure Legends

**Supplementary Figure (A)** Representative M-mode echocardiography of the left ventricle (LV) of the heart of young (3M) and old (18M) mice. **(B)** Ejection fraction (%) of the heart of young (3M) and old (18M) mice (n=6 group). **(C)** Adiponectin level (μg/ml) of the LV of the heart of young (3M) and old (18M) mice (n=6 group). **(D)** Leptin level (μg/ml) of the LV of the heart of young (3M) and old (18M) mice (n=6 group).

